# Supercontinuum intrinsic fluorescence imaging heralds ‘free view’ of living systems

**DOI:** 10.1101/2024.01.26.577383

**Authors:** Geng Wang, Lianhuang Li, Xiaoxia Liao, Shu Wang, Jennifer Mitchell, R.A.C “Chanaka” Rabel, Shirui Luo, Jindou Shi, Janet E. Sorrells, Rishyashring R. Iyer, Edita Aksamitiene, Carlos A. Renteria, Eric J. Chaney, Derek J. Milner, Matthew B. Wheeler, Martha U. Gillette, Alexander Schwing, Jianxin Chen, Haohua Tu

## Abstract

Optimal imaging strategies remain underdeveloped to maximize information for fluorescence microscopy while minimizing the harm to fragile living systems. Taking hint from the supercontinuum generation in ultrafast laser physics, we generated ‘supercontinuum’ fluorescence from untreated unlabeled live samples before nonlinear photodamage onset. Our imaging achieved high-content cell phenotyping and tissue histology, identified bovine embryo polarization, quantified aging-related stress across cell types and species, demystified embryogenesis before and after implantation, sensed drug cytotoxicity in real-time, scanned brain area for targeted patching, optimized machine learning to track small moving organisms, induced two-photon phototropism of leaf chloroplasts under two-photon photosynthesis, unraveled microscopic origin of autumn colors, and interrogated intestinal microbiome. The results enable a facility-type microscope to freely explore vital molecular biology across life sciences.

## Introduction

Focusing picosecond-femtosecond optical pulses on a nonlinear medium can generate supercontinuum signal highly informative of the medium itself (*1*). It raises the interesting possibility of comprehensive (spectroscopic) optical biopsy (*2*) or other applications of femtosecond biophotonics (*3*) if the often-studied transparent bulk gases/solids are replaced with live biological samples as the nonlinear media. However, the generated supercontinuum signals of plasma luminescence typically from picosecond irradiation (*4*) and white-light filamentation typically from loose femtosecond focusing (*5*) have been accompanied by optical breakdown (photodamage), which has been harnessed for precision surgery and micromachining via tight femtosecond focusing (*4, 6*). A lowered irradiance on the biological samples, intended for high resolution label-free multiphoton imaging (*7*), suppresses this supercontinuum generation around the near-infrared incident band (*4*) but often induces new molecules (photodamage) that emit supercontinuum-like (blue-to-red) fluorescence, i.e., ‘white’ flashes (*8*). Thus, hitherto observed biological supercontinuum signals have been universally associated with various photodamages detrimental to sample health and artifact-free image analysis (*9*). It is thus a significant finding that living samples can tolerate rather high irradiance up to one-half of water optical breakdown before nonlinear photodamage onset, if successive illumination pulses are spatially separated by ~1 diffraction-limited resolution (*10*). Here, within this narrow window of illumination, we find that supercontinuum (340-740 nm) fluorescence can be safely generated by a 1110-nm excitation to provide rich biological information, via the classic mechanism discovered by Göppert-Mayer (*11*) in 1931 but in a special form of simultaneous two-, three-, and four-photon absorptions of *intrinsic fluorophores* (*2*).

We began with simultaneous label-free autofluorescence-multiharmonic (SLAM) microscopy capable of imaging cellular flavin adenine dinucleotide (FAD) and reduced nicotinamide adenine dinucleotide (NADH) via two- and three-photon excited intrinsic fluorescence (2PIF and 3PIF), respectively, by a single-shot of filtered supercontinuum excitation (*12*). Given the same excitation band, we asked whether tryptophan (*13, 14*) could be simultaneously visualized via four-photon excited intrinsic fluorescence (4PIF), and whether two-photon excited long-wavelength intrinsic fluorescence (2PLIF) to the red of 2PIF could yield independent contrast from porphyrin- and/or lipofuscin-like biomolecules (*2*). The envisioned 4-color intrinsic fluorescence would thus benefit from the photon order super-multiplexed imaging (*15*) and common photon-order multicolor imaging (*16, 17*), but in a label-free manner (Fig. 1A). Due to plausible weak intrinsic fluorescence in the short-wavelength end, we reengineered the laser source (*18*) to balance various multiphoton processes (Fig. 1A) without inducing nonlinear photodamage (*10*). The resulting ‘supercontinuum’ intrinsic fluorescence imaging (SCIFI) naturally included fluorescence lifetime imaging microscopy (FLIM) (*19*) and two additional colors of epi-detected second- and third-harmonic generation microscopy (SHG and THG) (*12*) as ‘free’ capabilities to yield a 6-color simultaneously acquired (co-registered) signal (Fig. 1B, fig. S1; table S1).

**Fig. 1.**
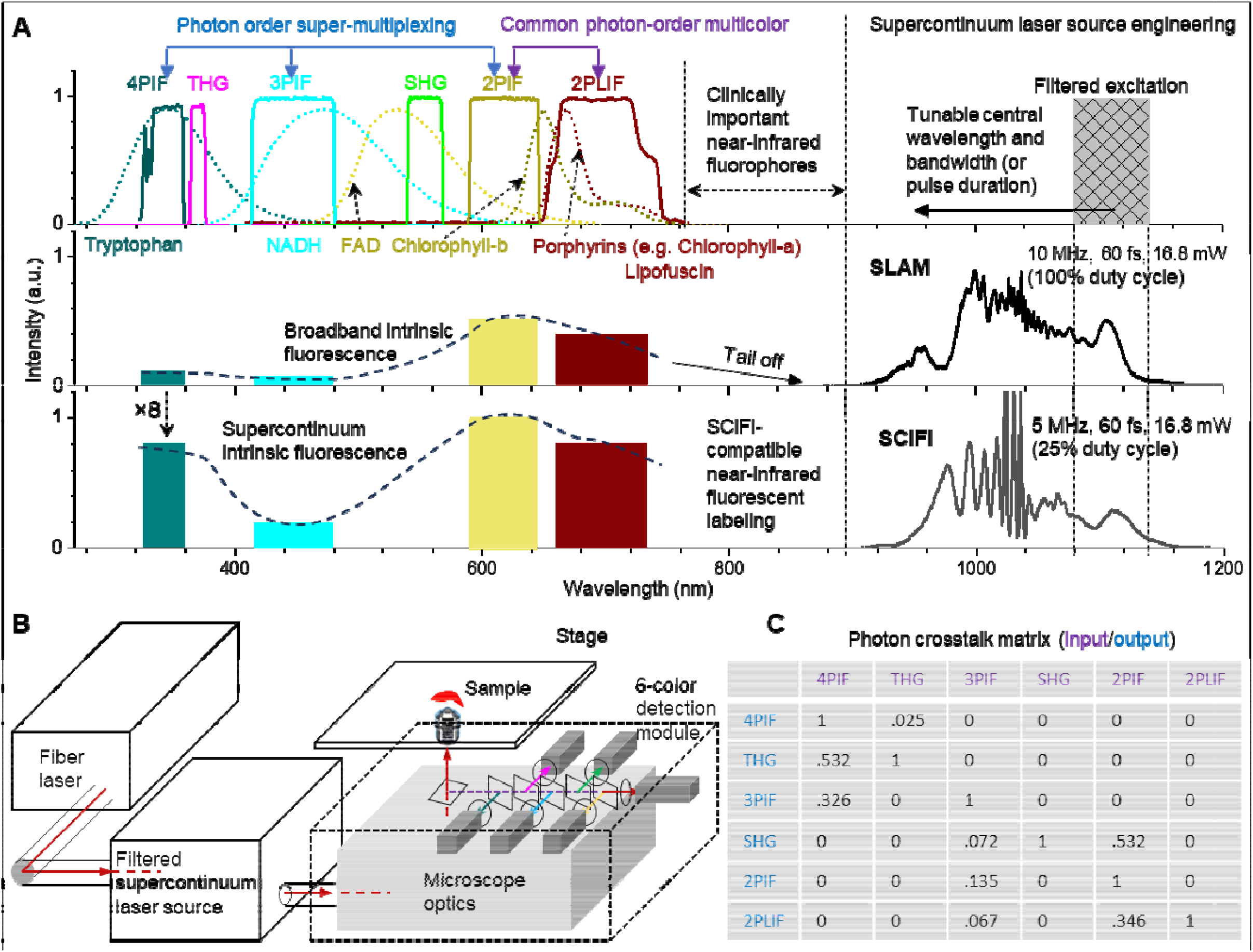
Concept and schematic of SCIFI. (**A**) Reported excitation in SLAM microscopy generates broadband intrinsic fluorescence from bandpass filter-separated cellular fluorophores (colored curves) weak at the short-wavelength end, whereas optimized excitation in SCIFI with reengineered supercontinuum source generates supercontinuum intrinsic fluorescence strong at the short-wavelength end, with 4× (or 2×) increased 4PIF/tryptophan (or 3PIF/NADH) relative to 2PIF/FAD due to 2× lower pulse repetition rate. Optional near-infrared fluorescent labeling extends the signal at the long wavelength end. (**B**) Schematic of an inverted laser-scanning microscope with automated stage and 6-color detection module. See fig. S1 for more details. (**C**) Photon crosstalk matrix of 6 detection colors from fluorophore solution-based calibration.

## Results

### Qualitative and quantitative live-cell SCIFI of animal samples

To appreciate qualitative SCIFI, we distinguished 4 stromal cell phenotypes in a niche of rabbit intestinal mucus (*20*) using 4PIF, 3PIF, and 2PLIF (Fig. 2A, left and middle; figs. S2-S4). The 2PLIF contrast across nucleus and cytoplasm, rather than 2PIF, allowed clear separation of two closely related cell phenotypes (Fig. 2A, middle). Similarly, independent 2PLIF contrasts were obtained in other samples to highlight specific cells (Fig. 2B, arrow) and extracellular components (Fig. 2B, arrowheads). It is therefore interesting to investigate whether these 2PLIF contrasts can be attributed to porphyrin(s) (*2*) or other intrinsic fluorophores suggestive of the embryogenesis-mimicking tumorigenesis (*21*) (fig. S5) and high-risk heart attack (*22*) (fig. S6). Moreover, we examined a pre-implantation *in vitro* fertilized bovine embryo near the 16-cell stage, coincident with the first lineage segregation into either the trophectoderm toward extraembryonic tissue or the inner cell mass toward the embryo proper (*23*). Remarkably, the observed 13 blastomeres (movie S1) were cleanly divided into 3 2PIF-visible versus 9 4PIF-visible blastomeres (Fig. 2C, left; fig. S7). As opposed to the largely even cytoplasmic distribution of differential 4PIF contrast over THG in the 4PIF-visible blastomeres near the 2-cell stage (movie S1, fig. S8), the corresponding uneven distribution near the 16-cell stage echoed the emergence of symmetry-breaking apical polarity cap at the 8-cell stage (*24*) (Fig. 2C, middle). These observations indicate the trophectoderm fate/lineage (4PIF-visible) and the inner cell mass fate/lineage (2PIF-visible) blastomeres near the 16-cell stage and the potential for clinically permissible label-free detection of embryo polarization without deep learning (*25*). It should be noted that the apical polarity cap is only detectable by 4PIF (or tryptophan).

**Fig. 2.**
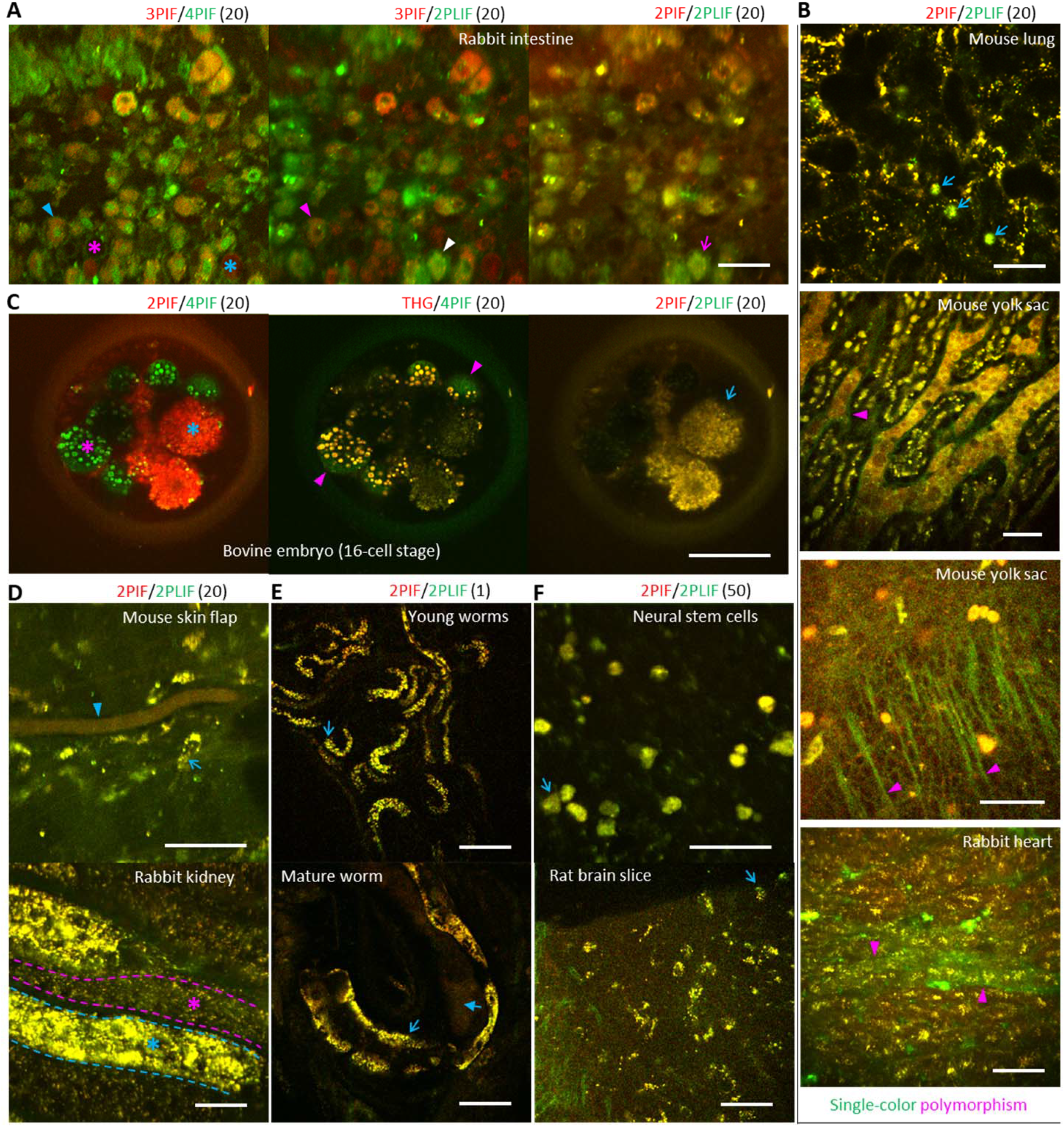
Visualization of live cells and extracellular matrix. The upper right corner of each image indicates detection color(s) and number of frame averaging. Scale bar, 50 µm. (**A**) Diverse stromal cells in a niche of rabbit small intestine with distinct color profiles including 4PIF and 2PLIF (stars, arrowheads, arrow). See figs. S2, S3 for more details. (**B**) Unique capability of 2PLIF to reveal stromal cells in mouse lung, emergent endothelial layer in yolk sac of one E11 mouse embryo (see fig. S5 for large-scale view), fiber-like structures in a vasculogenic field of E11 mouse embryo surface, and plausible atherosclerotic plaque in rabbit heart (see fig. S6 for more details). (**C**) Unique capability of 4PIF to reveal two distinct cell/blastomere phenotypes (stars), two blastomeres with apical polarity cap and the trophectoderm fate (arrowheads) in contrast to another blastomere with the inner cell mass fate (arrow) in a 16-cell-stage bovine embryo. See fig. S7 and movie S1 for more details. (**D**) Blood (arrowhead) versus stromal cells (arrow) in mouse skin flap and epithelial cells in rabbit kidney with two metabolic states (stars). (**E**) cells in the soma (arrows) or germline part (star) of young/mature *C. elegans*. (**F**) Cultured neural stem cells versus neurons in an area of rat brain slice near suprachiasmatic nucleus (arrows).

To demonstrate quantitative imaging (*19*), we scanned diverse samples with segmented cells either *in vitro* or *in situ* (*ex vivo* or *in vivo*) at a similar depth of 10-20 µm (tables S2-S4; figs. S3, S9). In contrast to the intestinal cells and other rare cells with distinct whole-cell 2PLIF (green contrast from porphyrin-like biomolecules) against 2PIF (Fig. 2A, 2B, arrows), most observable cell types have overlapping 2PIF and 2PLIF contrasts from orangish to greenish yellow in the form of punctuated intracellular granules (Fig. 2C-2F, arrows) and correlated 2PIF-2PLIF signal strengths across samples (Fig. 3A, fig. S10A), consistent with localized FAD and lipofuscin respectively in mitochondria (*13*) and lysosomes (*26*) according to their emission spectra (*2*). Assuming a tryptophan-NADH-FAD-lipofuscin model of animal cells with solution-based calibration by a photon crosstalk matrix (*19*) (Fig. 1B, right), we quantified the dimensionless extent of cellular lipofuscin by the ratio of crosstalk-compensated 2PLIF over pre-compensated 2PLIF, i.e., lipofuscin fluorescence independence extent (LFI) (Fig. 3B, top). This tetra-fluorophore cell model and related compensation was validated by the selective clearance of SHG signal in cells free of extracellular collagen (fig. S3) but not in those known to mingle with the collagen (Fig. 3C, arrowed segments; fig. S11).

**Fig. 3.**
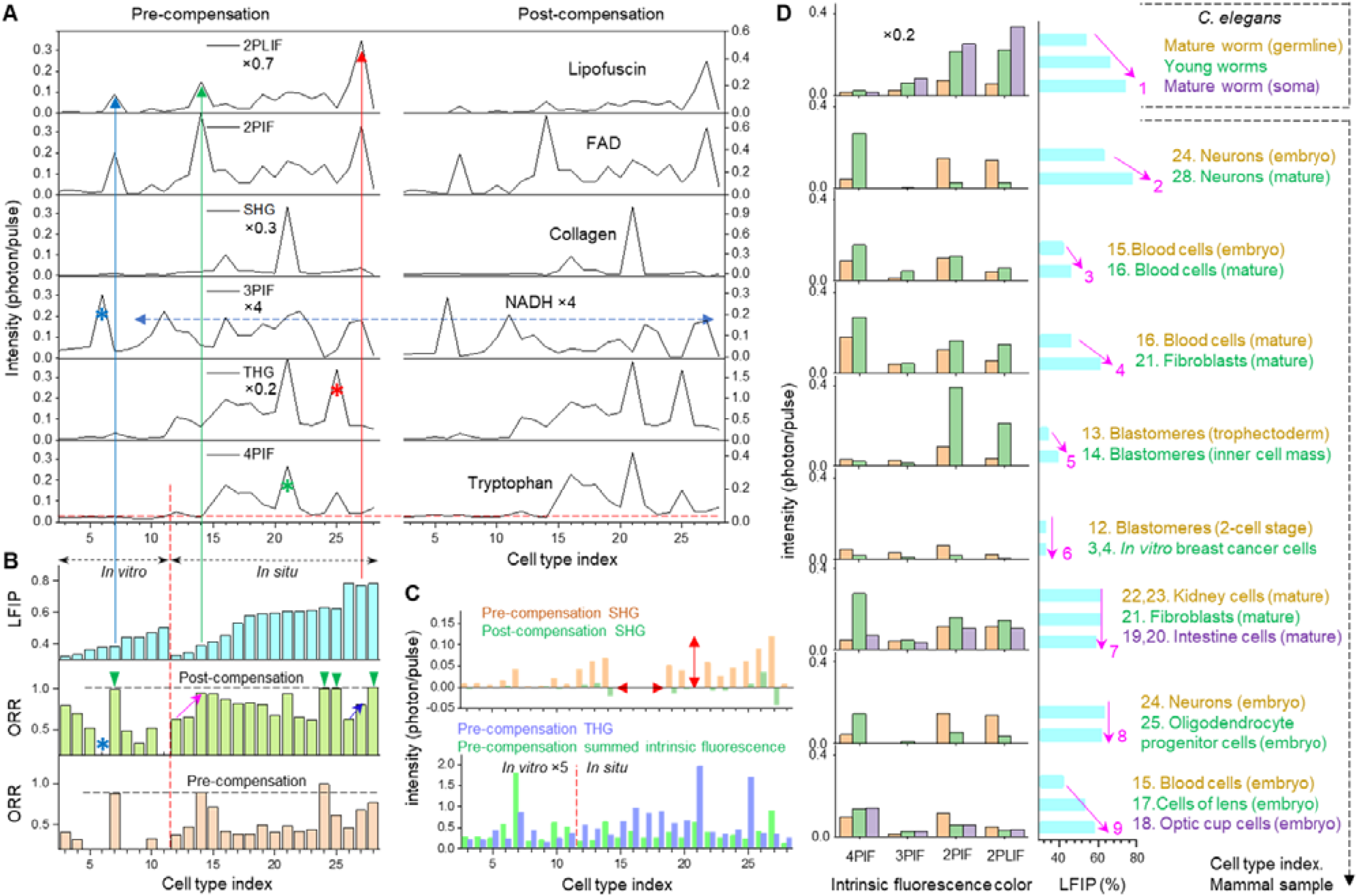
Quantitative imaging across cell types and samples. (**A**) Observed SCIFI 6-color profiles for 28 cell types from segmented individual cells in different fields-of-view of different samples (table S2). The signals may be scaled for clarity before (left) and after crosstalk compensation (right). (**B**) Quantified dimensionless LFI (top) and ORR before (middle) and after the crosstalk compensation (bottom) for the 28 cell types. (**C**) SHG clearance before and after the crosstalk compensation for most of the 28 cell types free of interference from collagen (top), and comparison of THG and summed intrinsic fluorescence intensities for *in vitro* versus *in situ* cell types before the compensation (bottom). (**D**) Correlation of observed LFI with cell aging or proliferation in diverse cell types (right) despite their highly irregular profiles of 4-color intrinsic fluorescence (left).

Taking this interference into account, we uniquely identified the fibroblasts cells in a collagen-rich microenvironment of mouse skin flap (Fig. 2D) by their peaked 4PIF across all observed cell types (Fig. 3A, green star). Similarly, the oligodendrocyte progenitor cells in a developing mouse brain (see below) and *in vitro* hamster kidney cells treated with staurosporine (STS) were uniquely identified by their peaked THG and 3PIF (NADH), respectively (Fig. 3A, red and blue stars). The latter is a known label-free biomarker of apoptosis (*27*) that may be extended to *in situ* situations. Among the three cell types with peaked 2PIF/2PLIF (Fig. 3A, arrows), the high LFI uniquely identified the epithelial cells in rabbit kidney (Fig. 3A-B, red arrow).

Surprisingly, high 4PIF intensities comparable to those of 2PIF/2PLIF were obtained across representative mammal cells, justifying the term of ‘supercontinuum intrinsic fluorescence’ despite ~4-time weaker 3PIF in between (Figs. 1A, 3A). The dynamic range of all colors were rather high (5×10^−4^ to 2 photons/pulse) so that specially calibrated analog photodetection (*19*) was developed to attain a photon-resolved quantification (Fig. 3A) beyond photon-counting, which is typically limited to a low throughput of <0.1 photon/pulse.

### Cellular biomarkers from aging-like stress to redox metabolism

To further understand LFI from the perspective of aging (*28*), we compared young vs. mature worms of *C. elegans* (Fig. 2E) and observed that the LFI of the former lies between that of the soma and germline part of the latter (Fig. 3D, Arrow 1). Also, we compared the mature brain of a rat (Fig. 2F; Fig. 4A, left) and the developing brain of a mouse embryo (Fig. 4A, right). In a myelination-induced ‘phase change’ during brain development detectable by SCIFI (Fig. 4A, 2PIF/4PIF images; movie S2), migratory oligodendrocyte progenitor cells (*29*) were easily discriminated against neurons (Fig. 4A, right). Observed LFI of the neurons in the mature brain was higher than that in the developing brain (Fig. 3D, Arrow 2). Moreover, the observed LFI of the red blood cells in skin flap of a mature (8-week-old) mouse (Fig. 2D, arrowhead) was higher than that in the developing eye of a mouse embryo (Fig. 4B, cyan arrowhead; Fig. 3D, Arrow 3). These observations strongly suggest a link between LFI and cell aging via a dimensionless extent of lipofuscin, which would avoid the poor specificity of lipofuscin fluorescence intensity across different samples (*30*).

**Fig. 4.**
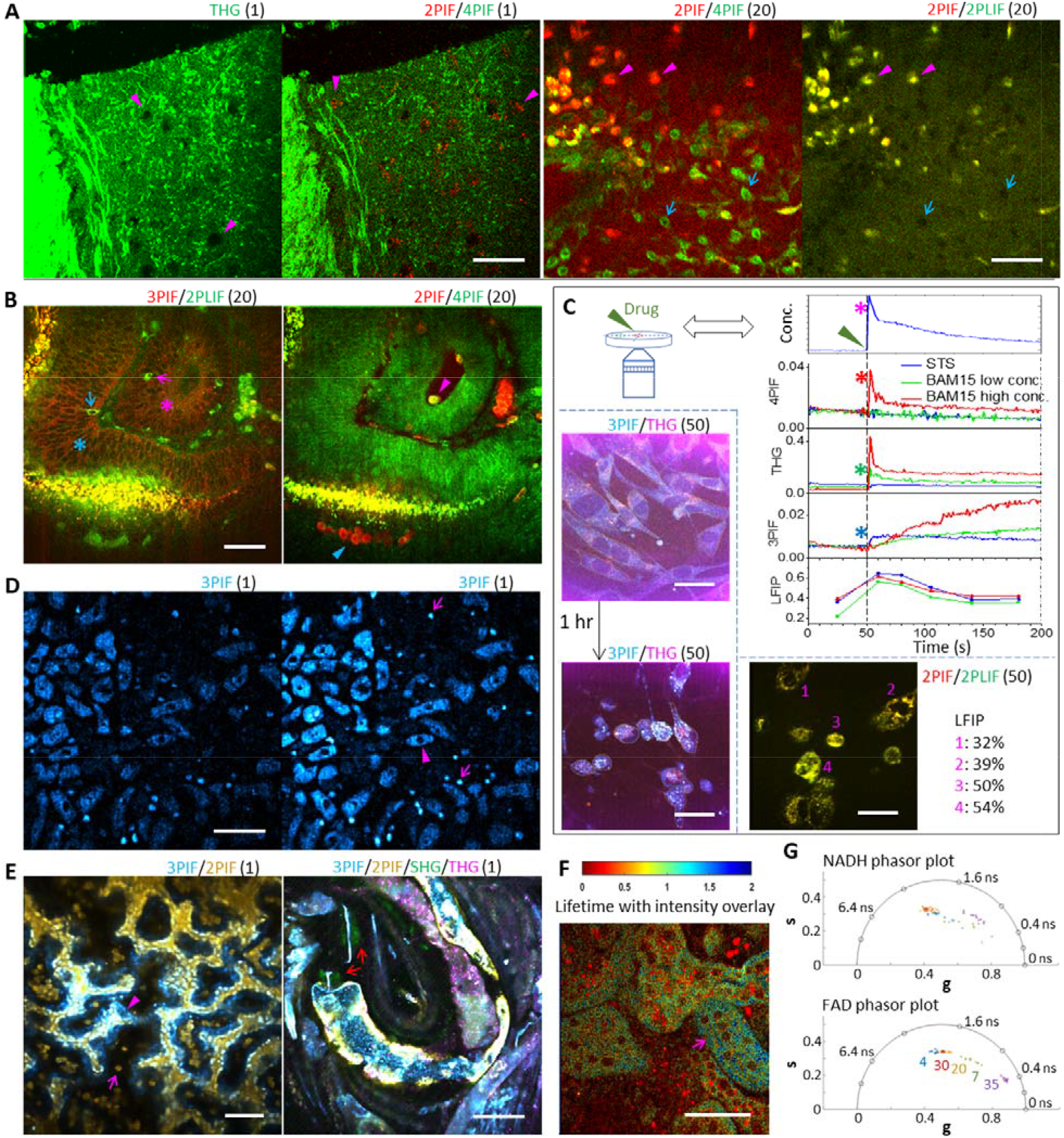
Versatile applications in biology and medicine. Scale bar, 50 µm. (**A**) Real-time (0.33 s per frame) 2PIF/4PIF imaging of brain slice (suprachiasmatic nucleus) of a 4-week-old rat reveals 2PIF-visible neurons (arrows) not obvious in THG imaging where the somata appear as dark shadows (arrowheads) in a continuous phase of myelin, and would thus enable more versatile targeted patching (left), while the related intrinsic fluorescence imaging of the *ex vivo* brain of an E15 mouse embryo reveals neurons (arrowheads) and oligodendrocyte progenitor cells (arrows) in a different state of continuous versus dispersive phases (right). See movie S2 in 3D for more details. (**B**) Developing ocular lens of an E11 mouse embryo and hosting optic-cup lined with similar fiber-like cells (stars) that contain rare cells (arrows and arrowheads). Refer to fig. S13 and movie S3 for more details. (**C**) Single-cell pharmacological imaging of cultured hamster kidney cells reveals temporal responses to externally introduced AP3, BAM15, and STS (colored curves) with *in-situ* monitored concentration (magenta star) detectable by AP3 fluorescence (top; see figs. S16, S17 and movie S4 for more details), conventional endpoint imaging of the cells before and after adding STS in different fields of view shows apoptosis-induced morphological change (bottom left) absent from AP3 exposure (see figs. S14, S15 for more details), and single-cell LFI analysis differentiates low-stress amoeba-shaped adipose-derived stem cells (1 and 2) from high-stress spherically shaped counterparts (3 and 4) (bottom right). (**D**) Time-lapse increase of 3PIF in rabbit intestinal cells (arrowheads) and microbes (arrows) as indicator for photodamage. See fig. S19 and movie S5 for more details. (**E**) UDVD denoising of real-time low-signal-to-noise raw video reveals 3PIF-visible cells of plausible endothelial origin (arrowhead) and a flowing blood cell (magenta arrow) in yolk sac from an E11 mouse embryo (left), and SHG-visible pharynx bulbs (red arrows) of a freely moving *C. elegans* (right). See figs. S20, S21 and movies S6, S7 for more details. (**F**) FLIM analysis of epithelial cells in mouse kidney reveals an unusual lobe (arrow) with long 4PIF lifetime (see fig. S24 for more details). (**G**) FLIM phasor plots of 3PIF/NADH (top and 2PIF/FAD (bottom) that show the better performance of the latter to discriminate numerous individual cells with related cell type indices (see table S2).

We observed an increased LFI from the blood cells in the skin flap to the surrounding less proliferate fibroblasts (Fig. 2D, arrowhead vs. arrow; Fig. 3D, Arrow 4). Also, an increased LFI was observed from the 16-cell stage bovine blastomeres with the trophectoderm fate to those with the inner cell mass fate (Fig. 3D, Arrow 5). Moreover, the lowest LFI (33%) was approached by bovine blastomeres near the 2-cell stage and an immortal cell line of breast cancer (Fig. 3D, Arrow 6). These observations seemed to correlate low LFI with high proliferation potential after different differentiated cell types experience different extents of aging-like stress to lose proliferation potential. Specifically, the epithelial (intestine/kidney) cells in the same 8-week-old mouse (fig. S9) experienced a similar stress as the fibroblasts to attain a comparable LFI (Fig. 3D, Arrow 7), whereas the neurons and oligodendrocyte progenitor cells in the mouse embryo experienced an accelerated stress to reach this LFI earlier (Fig. 3D, Arrow 8). The increased LFI (Fig. 3D, Arrow 9) from the fiber-like cells lining the ocular lens of the developing eye (Fig. 4B, magenta star) to those lining the hosting optic cup (Fig. 4B, cyan star) supported the paracrine signaling from a preexisting optic cup to a lately developed more proliferate lens (*31*), leading to a converged metabolism (6-color profile) and morphology of these cells (figs. S12) despite the rather different origins of the optic cup and lens (*32*).

In contrast to the other colors, the crosstalk compensation significantly modified the weak 3PIF (Fig. 3A, arrowed segment) and thus enabled accurate metabolism-based quantification of optical redox ratio (ORR) (*26, 33*). We observed that rabbit kidney cells in proximal versus distal tubules (*16*) (Fig. 2D, stars) with rather different ORRs (Fig. 3B, blue arrow) exhibited a largely converged LFI (77-78%) or proliferation. This effect is in sharp contrast to the Warburg effect that enhances cell proliferation by a shifted metabolism (*34*). As another example, the blastomeres with the inner cell mass fate near the 16-cell stage (ORR 0.95) emerged from the blastomeres near the 2-cell stage (Fig. 3B, magenta arrow) that approximated the blastomeres with the trophectoderm fate near the 16-cell stage (ORR 0.66), despite the rather similar LFI among the three (34-39%). With similarly strong FAD signal, the mutated neural stem cells in this study and reported epithelial cancer stem cells (*35*) might mimic the high-ORR blastomeres (Fig. 3A-B, blue and green arrows) for immortality and tumorigenesis (*21*), respectively. On the other hand, near unity ORR was approached by *in vitro* neural stem cells (Fig. 2F), the neurons and oligodendrocyte progenitor cells in the mouse embryo, and the neurons in a mature rat, despite their rather different LFI across 38-78% (Fig. 3B, green arrowheads). This high ORR discriminated the *in vitro* neural stem cells from the bovine blastomeres with the inner cell mass fate (Fig. 3A-B, blue vs, green arrows) to highlight the unique metabolism of brain cells, which would be missed without the crosstalk compensation.

Taken together, the observed LFI in this study and the well-known ORR are different dimensionless cellular biomarkers complementary with each other. They comprehensively discriminated rare cells from their neighboring cells in the developing eye (fig. S13; movie S3) not possible using conventional histology and phase-contrast microscopy (*32*).

### Multiparametric endpoint and kinetic in vitro assays

We then evaluated the interaction between *in vitro* hamster kidney cells with a drug or imaging agent (table S2). First, the treatment by STS induced apoptosis via a modified cell morphology, which was detected by an SCIFI endpoint (or snapshot) assay involving different fields (Fig. 4C, bottom left). In comparison to the treatment-free control, these apoptotic cells retained a low LFI despite a dramatically reduced ORR (Fig. 3B, blue star), suggesting the independence between LFI and apoptosis. Second, a treatment by the weight-loss drug BAM15 (*36*) likely generated an aging-like stress with increased LFI (44% versus 32%/control), even though the 6-color profiles of treated and untreated cells were largely comparable (table S4). Third, using a membrane dye designed for SHG imaging with low fluorescence background (AP3) (*37*), the corresponding endpoint assay validated the SHG imaging of cell membrane but detected a significant increase of 2PIF/2PLIF signals (fig. S14), which was confirmed using the breast cancer cells (fig. S15) and called for a detailed examination. In summary, the endpoint assay is limited by the different fields of view that prohibit a more rigorous understanding.

To overcome this limitation, we employed the time-lapse or kinetic assay to track cellular changes in the same field of view (*33, 38*) (Fig. 4C, top right). The high sensitivity of SCIFI detected the unexpectedly strong AP3 fluorescence via 2PLF/2PLIF from the empty spaces among the cells, enabling *in-situ* monitoring of concentration profile (Fig. 4C, magenta star). Thus, the increased 2PIF/2PLIF signals in the endpoint and kinetic assays can be attributed to the surprising ability of AP3 to fluorescently label the cytoplasm (figs. S14, S15). At a high concentration, 4PIF ‘sensed’ cytotoxicity in real-time plausibly due to the inhibited Förster resonance energy transfer (*14*) by AP3 (fig. S16), which was confirmed by BAM15 (Fig. 4C, red star). We thus limited the maximal concentration of this transient exposure to avoid the resulting cytotoxicity (fig. S17), allowing the cells to instead ‘sense’ BAM15 undergoing endocytosis via THG-visible nanovesicles (*39*) (Fig. 4C, green star; movie S4). In contrast, these cells responded to apoptosis-inducing staurosporine (STS) via 3PIF/NADH (*27*) (Fig. 4C, blue star) but not THG (fig. S17), suggesting a non-nanovesicle entry for STS. Regardless of the mode of toxic action, potential drugs (or SCIFI-compatible labeling fluorophores in Fig. 1A) should attain efficacy (or effectiveness) below a concentration threshold (*40*) to avoid any SCIFI detectable cytotoxicity.

Our real-time assay of LFI also revealed an agile cellular recovery from the transient exposure to STS or BAM15 not observable from 2PIF/2PLIF intensities (Fig. 4C, red curves; fig. S17). The recovery preceded the plausible morphological change of STS-induced apoptosis observed from the endpoint assay (compare fig. S17C with Fig. 4C, bottom left). This effect would be challenging to detect from the endpoint assay due to the inaccurately quantified LFI of hamster kidney cells with intrinsically low 2PLIF (table S4), highlighting a unique advantage of the kinetic assay. On the other hand, our LFI-based endpoint assay may serve as a general label-free assay (*41*) for stress test of custom-cultured cells (Fig. 4C, bottom right) independent of the NADH apoptosis biomarker or other cell death assays. This endpoint assay produced low and relatively constant 4PIF (0.015-0.027 photon/pulse) for 4 different *in vitro* cell types without or with drug treatments, which may be used as a biomarker to discriminate *in situ* against *in vitro* cell types (Fig. 3A, broken lines). To improve drug discovery, realistic 3D disease models such as organoids should approximate the pertinent *in situ* cell types in 4PIF/THG and summed intrinsic fluorescence intensities, which are typically ~4-time higher than those of *in vitro* cell types (Fig. 3C, bottom).

### Performance-enhancing capabilities at minimum cost

First, using broadband ‘hyper-fluorescence’ as an intrinsic indicator of photodamage (*10*), we routinely collected strong 4PIF and 3PIF without compromising sample health (fig. S18). However, 10% increased power beyond our typical imaging condition (table S1) induced uniform hyper-fluorescence to the *in-situ* cells of rabbit small intestine (Fig. 4D, fig. S19), but not the intracellular ‘hot spots’ inside *in vitro* cells (*8, 10*) (movie S5). This is worth further study because primary cells are typically more heterogeneous than cell lines.

Second, we compared three self-supervised deep learning models in ‘blind-spot’ video denoising free of ground truth data (*42-44*). The overall high performance of UDVD denoising (*44*) under both fast and slow sample motion (figs. S20, S21; movies S6, S7) may mitigate the low signal-to-noise ratio intrinsic to fast ‘gentle’ imaging (Fig. 4E). In a real-time fashion (*43*), UDVD may enable intraoperative assessment in surgical oncology (*45*). Also, the stripe self-correction of mosaicking-assisted image acquisition (*46*) helps reveal a novel large-scale structure of rat suprachiasmatic nucleus (figs. S22, S23) while 2PIF/4PIF visualization can improve targeted patching in 3D over THG triage (*47*) (Fig. 4A, left; movie S2).

Third, the built-in FLIM further quantified fluorescence for intestinal cell phenotyping (fig. S3) and distinguished individual cells of different cell types by 2PIF/FAD (Fig. 4F; figs. S24-S26). It may be tempting to supersede SHG/THG with FLIM-empowered intrinsic fluorescence. This is not justified as SHG has the highest sensitivity to detect extracellular collagen fibers (fig. S24) and extrinsic food starch particles (fig. S25), whereas the anticipated collagen fluorescence (*2*) across 4PIF-3PIF spectral range is universally negligible (e.g., see fig. S27). Also, THG complements intrinsic fluorescence to identify intracellular vesicles and possibly extravasated red blood cells in *ex vivo* rabbit kidney (fig. S27). The THG intensity in ~10-nm bandwidth is comparable to summed intrinsic fluorescence intensity of 4PIF, 3PIF, 2PIF, and 2PLIF (Fig. 3C, bottom), and may be the most sensitive imaging for photodamage-prone samples at an attenuated illumination. Somewhat unintuitively, the apparent correlation in signal strength from the THG-4PIF or 2PIF-2PLIF pair (Fig. 3A) does not prevent independent information from the two colors (table S5).

### Applicability in plant leaves under photosynthesis

We subsequently tested the utility of SCIFI beyond animal kingdom. From a detached autumn common (evergreen) ivy leaf, the 2PIF-revealed chloroplasts likely by chlorophyll-b fluorescence (*48*) (Fig. 1A) are cleanly separated from 3PIF-revealed vacuoles of palisade mesophyll cells by stilbene (*49*) and from 2PLIF-revealed photosystem II (PSII) structures by intense and thus 20×-attenuated chlorophyll-a fluorescence (Fig. 5A, table S6). The PSII structures at the peripherals of the vacuoles are identified by the Kautsky effect (*50*) (Fig. 1A, right; movie S8). This observation defies the conventional wisdom of spatially restricted chlorophyll-a to chloroplasts and extends conventional visible/single-photon photosynthesis (*51*) to near-infrared/two-photon photosynthesis, which resembles the observation of human two-photon vision (*52*) and emerges as an alternative to study the ultrafast phenomena of ambient photosynthesis using exotic optical source and detection (*53*). Meanwhile, a two-photon counterpart of chloroplast phototropism (*54*) may lead to unusual chloroplast movement and emergence (Fig. 5B, fig. S28; movies S8, S9). Without the interference of this effect in one cycle of the Kautsky effect, the chloroplasts exhibit *increasing* 2PIF (Fig. 5A, right; movie S8), a subtle accompanying effect that has been masked by the intense but *decreasing* chlorophyll-a fluorescence from standard measurements (*51*). This dynamic effect from chloroplasts also occurs at up to 50% weaker illumination powers (table S1) with a refractory period within 10-20 min and a dark acclimation time of >60 min (fig. S29), and thus reflects the physiological ‘vitality’ of the chloroplasts rather than a photodamage effect. As to static analysis, a yellow/diseased ivy leaf can be differentiated by 3 SCIFI visibility markers related to the chloroplasts, vacuoles, and PSII structures (Fig. 5C, table S6). To correlate this pathological coloration in ivy leaves with the autumnal coloration in maple leaves, we employed these markers to differentiate major leaf phenotypes (table S6), including those from an autumnally variegated green-yellow-red maple leaf (Fig. 5D) that enables direct comparison of different colored sectors in the same leaf (*55*).

**Fig. 5.**
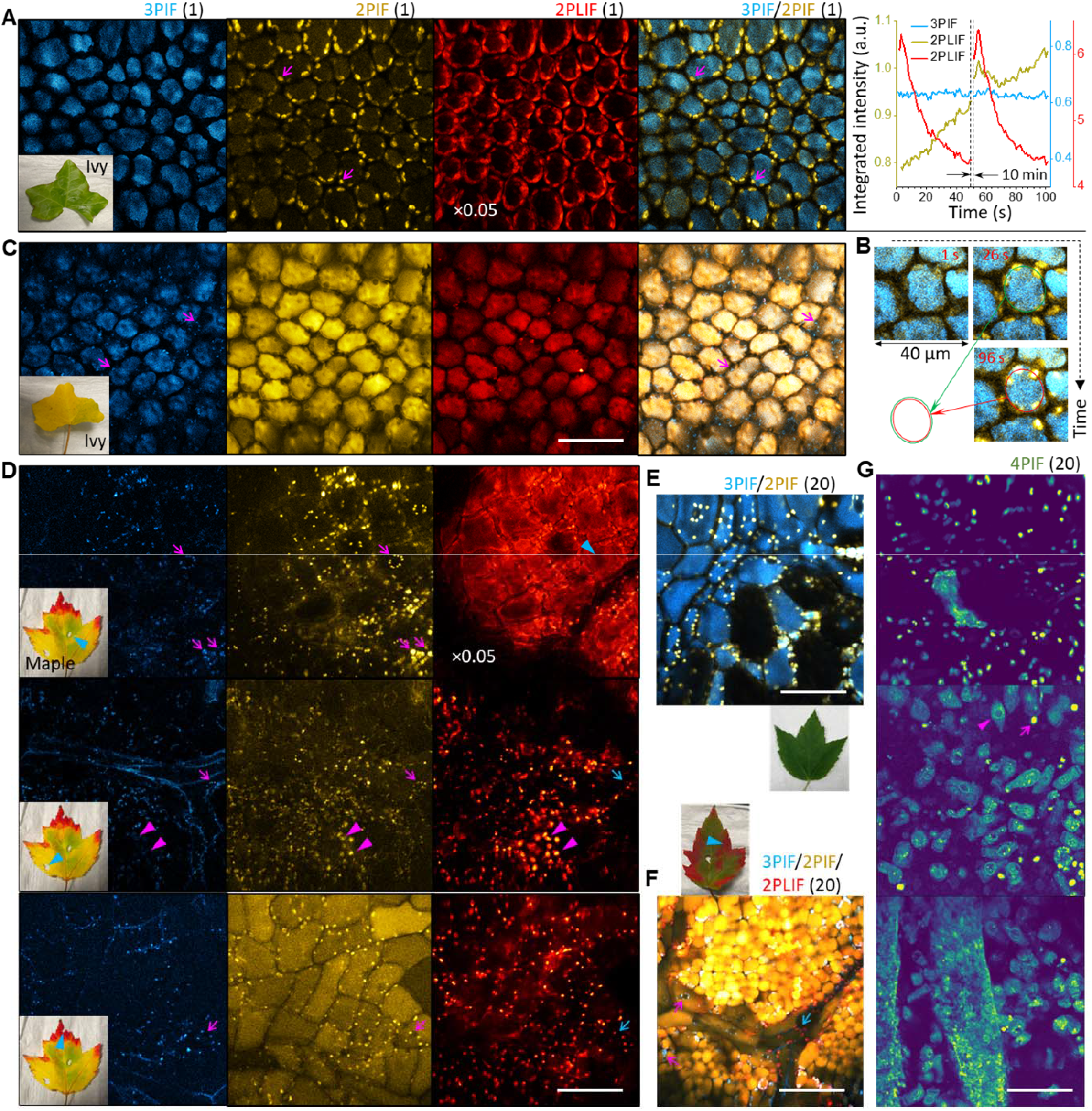
Label-free imaging of plants and microbes. Scale bar, 50 µm. (**A**) Green autumn ivy leaf with 3PIF-visible vacuoles, 2PIF-visible chloroplasts (arrows), and ×0.05-scaled 2PLIF-visible PSII structures undergoing recurrent Kautsky effect (red curves) with increased chloroplast visibility (yellow curves) but constant vacuole visibility (cyan curves). See figs. S28, S29 and movie S8 for more details. (**B**) Coordinated movement of chloroplasts toward each other in one mesophyll cell during imaging (from a green oval to a smaller red oval that links the chloroplasts). See fig. S28 and movie S8 for more details. (**C**) Yellow/diseased autumn ivy leaf with 3PIF-visible vacuoles and chloroplasts (arrows), 2PIF-visible vacuoles, unattenuated 2PLIF-visible vacuoles without observable PSII structures, and composite 3PIF/2PIF architecture opposite to that of its green counterpart. (**D**) Autumnally variegated green-yellow-red maple leaf showing 3PIF- and 2PIF-visible chloroplasts (magenta arrows) invisible among ×0.05-scaled 2PLIF-visible PSII structures (cyan arrowhead) but visible in unattenuated 2PLIF (cyan arrows) without observable PSII structures. The yellow part of this leaf has 2PIF-visible but 3PIF-invisible chloroplasts (magenta arrowheads) specific to a green spring maple leaf (see fig. S30 for more details). (**E**) Green spring maple leaf showing composite 3PIF/2PIF architecture like that of the green autumn ivy leaf. See fig. S30 for more details. (**F**) Autumnally variegated green-red maple leaf with 3PIF-visible only (magenta arrows) and 2PLIF-visible only chloroplasts (cyan arrows) in the red part. See fig. S31 for more details. (**G**) Depth-resolved detection of microbes (arrow) and/or stromal cells (arrowhead) from mucus (top) to epithelium (bottom) in rabbit small intestine. See fig. S33 for more details.

Our leaf phenotyping attributes the difference between green autumn ivy versus maple leaves to a seasonal change of the latter rather than a difference between evergreen versus deciduous species (Fig. 5A, 5E vs Fig. 5D, top; fig. S30). Prominent PSII structures mask the 2PLIF visibility of chloroplasts (Fig. 5D, top) and extend well beyond them (movie S8). Maple leaf autumnal red/yellow coloration is marked by disappeared PSII structures with 20×-attenuated chlorophyll-a fluorescence, which exposes the otherwise masked 2PLIF-visible chloroplasts (Fig. 5D, top vs. middle/bottom). The related autumnal red coloration emerges expectedly from *de novo* production of anthocyanin (*56*) that enables vacuole visibility (*49*) via 2PIF (Fig. 5D, bottom), resembling the altered vacuole visibility in the diseased yellow ivy leaf (Fig. 5C). However, the corresponding 3PIF- and 2PLIF-visible chloroplasts may develop into one type of ‘non-vital’ 3PIF-visible only chloroplasts present also in the yellow ivy leaf (compare Fig. 5F and Fig. 5C, magenta arrows) and another type of 2PLIF-visible only chloroplasts to be recycled before leaf fall (compare Fig. 5F and Fig. 5D bottom, cyan arrows; fig. S31). In contrast, maple autumnal yellow coloration emerges without the altered vacuole visibility (Fig. 5D, middle vs. top) in the ivy pathological yellow coloration (Fig. 5C), but with spring 2PIF-visible but 3PIF-invisible chloroplasts (Fig. 5E) absent from a green autumn maple leaf (Fig. 5D, top; fig. S30). Taken together, a yellow autumn leaf may use additional energy to compensate the lost ‘vitality’ in 3PIF-visible autumn chloroplasts by *reproducing* the ‘vital’ spring chloroplasts (movie S10), paralleling the *de novo* production of anthocyanin in a red autumn leaf (fig. S32). The underlying hypothesis suggests that autumnal yellow and red phenotypes emerge independently from autumnal green phenotype (table S6), i.e., no yellow state mediates between autumnal green and red states (*56*).

### Beyond eukaryotic cells and conventional multiphoton microscopy

Finally, we applied SCIFI to prokaryotic cells with relevance in the food industry (*41*). The microbes in rabbit intestinal mucus (fig. S33) exhibited abnormally high LFI (83%, table S4), which we attributed to the strong 2PLIF from a chlorophyll-a-rich diet (*57*) rather than an aging-like stress. To evaluate the possible diet origin of the abnormally strong 4PIF from these microbes, we examined diverse vegetation and identified strong or independent 4PIF only in rare cases, such as the leaf stalk of mimosa pudica and the wall of ginger oleoresin cells (figs. S34, S35; movie S11). Thus, the strong bacterial 4PIF arises more likely from intrinsically abundant tryptophan (*58*) than diet, whereas the advantage of imaging over traditional non-imaging spectroscopy may empower food quality assessment (*59*). The strong 4PIF signal allows real-time depth-resolved *in-situ* detection of intestinal microbes (Fig. 5G) potentially valuable in gut biopsy or intravital imaging (*20*), while combined 4PIF and THG uniquely reflects the evolution from the prokaryotic cells of intestinal microbes to the eukaryotic cells of *C. elegans* and then to the eukaryotic cells of mammals (fig. S10C). The absence of the microbes near intestinal epithelium and their abundance in cell-free mucus (Fig. 5G, bottom vs. top) were intermediated by a moderate presence with spatially separated stromal/immune cells (Fig. 5G, middle), suggesting the existence of a layered immunological barrier in mucus to explain the intrigue observation why most crypts were not colonized by bacteria (*60*). Interestingly, the microbes share the same photodamage dynamics as their host cells (Fig. 4D, fig. S19; movie S5), and can therefore be discriminated against non-living particles with no hyper-fluorescence.

To summarize, the intense chlorophyll-a fluorescence in plants and microbes implicates context-aware near-infrared fluorescently labeled imaging of animal samples that removes the restriction to label-free imaging and thus bridges preclinical/labeled and clinical/label-free studies (Fig. 1A, table S7). This imaging can be further empowered by replacing the dichroic/filter-based detection with an optical fiber-coupled spectroscopic detection (*61*) to expand the label-free molecular imaging (*2, 49*) beyond a few preselected targets (Fig. 1A). With a similar capital expense as conventional multiphoton microscopy (table S8) but a potentially lower operating expense (fig. S36A), SCIFI gains some advanced capabilities largely *free of cost* (table S9) and versatile adaptations *free of restriction* (table S10). In particular, the stabilized supercontinuum source and subsequent optical fiber-coupled delivery (*18*) not only simplifies routine maintenance by the laser-microscope alignment decoupling (*62*), but also enables tunable or multi-band excitation and simultaneous non-imaging applications such as optogenetics (*63*) by harnessing unused (unfiltered) portions of the supercontinuum (Fig. 1A). With available tools to democratize the imaging (fig. S36B) and further optimization of illumination condition (see supplementary text in the supplementary materials), SCIFI heralds ‘free view’ of living systems, especially those close to our everyday life.

## Author contributions

H.T. conceived the idea. G.W., L.L., X.L., J.C., and H.T. designed the materials and methods. G.W., L.L., J.M., R.A.C.R., and D.J.M. prepared the materials and conducted experiments. X.L., S.W., S.L., A.S., and J.C. developed software and/or performed image processing. G.W., L.L., X.L., M.B.W., M.U.G., J.C., and H.T. analysed the results. G.W., L.L., and X.L. drafted the manuscript. J.C. and H.T. edited the manuscript with input from all authors.

## Acknowledgments

We thank Jindou Shi, Janet E. Sorrells, Rishyashring R. Iyer, Edita Aksamitiene, Carlos A. Renteria, Eric J. Chaney, and Stephen A. Boppart for their general assistance during this study. This work was supported by grants from the National Institutes of Health, U.S. Department of Health and Human Services (R01 CA241618) and the Natural Science Foundation of China (Grant No. 81671730).

## Data and materials availability

All data are available in the main text or the supplementary materials.

